# Rearing and Head Scanning as Functionally Equivalent Information-Seeking Behaviors

**DOI:** 10.64898/2026.04.30.721974

**Authors:** Ryan Troha, Billy Gregory, Dylan Burks, Andrew Petro, Kaci Kirkpatrick, Ehren Newman

## Abstract

Spatial memory is crucial for navigation and adapting to changing environmental conditions. Known neurophysiological mechanisms of spatial memory center on the importance of hippocampal activity and its spatial tuning. Yet, the behavioral strategies that support adaptive spatial encoding remain poorly understood. We have shown that dorsal hippocampal activity during rearing is necessary for spatial working memory, highlighting a role of information seeking behaviors for spatial memory encoding. Similarly, spatial tuning by dorsal hippocampal neurons is substantially updated during another information seeking behavior: attentive head scanning. However, the functional relationship between these behaviors is unknown. Here, to assess the relevance of environmental context for the expression of these behaviors, we quantified rearing and head scanning in a radial-arm-maze spatial working memory task while manipulating the height of the maze walls. Our goal was to test whether the stereotyped patterns of rearing that rats generate with tall walls are replaced with attentive head scanning when the walls are short enough to reach the top without rearing. We found that rats reared significantly less often when the walls were shortened and, instead, exhibited frequent attentive head scanning. The head scanning was done when and where the rats had previously exhibited stereotyped rearing. These results support the hypothesis that rearing and head scanning are functionally related behaviors. Future work should test two key inferences: 1) Head scanning is a critical epoch of spatial memory encoding, and 2) Spatial tuning by hippocampal neurons is updated during rearing.

**Significance statement:** Spatial memory is a core cognitive function, essential for healthy independent living. Though the hippocampus is critical for spatial memory, it remains unclear when and how. Separate prior studies link rearing and lateral head scanning to key periods of hippocampal processing, suggesting both behaviors support sensory information gathering for updating cognitive maps. However, their relationship is unresolved. Here, we test whether these behaviors are functionally interchangeable, with environmental structure determining expression. In a radial-arm maze, rats reared frequently with 21 cm walls but showed reduced rearing when walls were shortened to 4.6 cm, instead increasing head scanning at similar locations. These findings suggest rearing and head scanning share underlying motivations and provide a basis for comparing hippocampal activity during exploration.

## Introduction

Spatial memory is a core cognitive competency, critical for effective navigation, adaptive behavior and healthy independent living. Understanding the mechanisms that support spatial memory is, thus, of high importance. The neurophysiological mechanisms supporting spatial memory have been well studied. The findings of which emphasize the importance of hippocampal activity and its spatial tuning. Yet, the behavioral contexts that engage these mechanisms to support spatial memory remain ill-defined. Tolman famously emphasized the importance of exploratory behaviors for spatial encoding (Tolman, 1948). Supporting this, we recently found that hippocampal activity when rats rear onto their hind legs is necessary for spatial working memory (Layfield et al., 2023). Separately, it has been shown that spatial tuning by hippocampal neurons is updated during another exploratory behavior: attentive head scanning (Monaco et al., 2014). The functional relationship between these exploratory behaviors is not well understood. Here, we report the findings of an experiment designed to test the hypothesis that rats treat attentive head scanning and rearing as interchangeable exploratory behaviors as a function of environmental context.

The pioneering work of Tolman demonstrated that intrinsically motivated exploration of an environment supports subsequent operantly motivated navigation. He suggested that this exploration enables the construction of a “cognitive map” (Tolman, 1948). Evidence of spatial tuning among neurons in the hippocampus led to the hypothesis that the hippocampus was central to the construction of the cognitive map (O’Keefe & Dostrovsky, 1971; O’Keefe & Nadel, 1978). This hypothesis that has been bolstered by further evidence of spatial tuning throughout the formation (Taube et al., 1990; Lever et al., 2009; Hafting et al., 2005; Sargolini et al., 2006; Taube 2007; Boccara et al., 2010) and demonstrations that impairments of hippocampal function are tied to deficits in spatial memory (Moser et al., 1993; Packard & McGaugh, 1996; Lee et al., 2019; Ferbinteanu et al., 2003). Yet, the link between hippocampal function and exploration-driven spatial learning remains poorly understood.

A major step forward towards linking exploration to hippocampal function came with the discovery that spatial tuning in the hippocampus was overtly updated during exploration (Monaco et al., 2014). While examining the spatial tuning of hippocampal CA1 neurons, Monaco et al. noted that a subset of neurons developed a new place field mid-session as rats completed laps on a circle track for food reward. They showed that, on the lap immediately prior to the first clear expression of a place field, the rat reliably stopped at the point where the field would emerge and performed attentive head scanning. That is, the rat paused and extended its head over the side of the track, which had low walls, and explored the context beyond the track boundary. This work demonstrated a connection between exploratory behavior and changes in hippocampal tuning but, critically, because there was no memory component to the task it did not address any putative connection to memory function.

Connecting hippocampal function and exploration-driven spatial learning directly, we recently found that selectively silencing dorsal hippocampal activity when rats reared onto their hind legs during the study phase of a spatial working memory task impaired their performance in a subsequent test phase (Layfield et al., 2023). Rats were performing a delayed spatial win-shift task on a radial arm maze, collecting food rewards from four open arms during a study phase and then, in a test phase following a delay of a few minutes, all eight arms were opened and additional reward pellets could be collected from the previously unvisited arms. Rats rear reliably and frequently in this task, often at the ends of arms after collecting rewards. This pattern of behavior seemed systematic and perhaps a spatial navigation strategy that rats used to complete the task, as has been suggested by previous work (Lever et al., 2006). However, no study has examined how rearing compares to other exploratory behaviors such as head scanning, which is known to be tied to hippocampal spatial processing (Monaco et al., 2014). We therefore sought to directly compare these behaviors in the same delayed win-shift task.

We hypothesize that attentive head scanning and rearing may similarly support hippocampal-dependent spatial encoding but that each behavior is motivated under different environmental contexts. Head scanning and rearing frequency both occur in novel testing environments - occurring less frequently with familiarization (Sutherland et al., 1982, 1983; Lever et al., 2006; Davis et al., 2025). Rearing occurs most frequently overall on the first trial in an open field and occurs less frequently over further subsequent trials, but then increases at the onset of each new trial (Lever et al., 2006). Likewise, head scanning occurs most often on the first trial on an elevated track and occurs less often over subsequent trials, but increases at the start of each trial (Davis et al., 2025). Notably, however, such examinations of when rats head scan or rear have focused on one or the other behavior. This may be because different environmental contexts (e.g., an open field surrounded by tall walls vs. an elevated track) differentially motivate one exploratory behavior over the other (e.g., rearing vs. head scanning).

In our prior work with the radial arm maze, we reliably observed rearing but not head scanning (unpublished data). The maze structure may be the reason. The maze had no top but the arms were enclosed by 21 cm tall clear walls. Thus, though the rats have visual access to the surrounding environment, they may have nonetheless been motivated to sample over the walls, requiring the rats to adopt a rearing posture. In contrast, the comparatively short circle track walls used by Monaco et al. (2014) may have required a less erect posture, resulting in apparent attentive head scanning. This prompts the question of whether head scanning and rearing are functionally similar exploratory behaviors, but occur depending on the structure of environmental boundaries.

Alternatively, differences in expression of rearing versus head scanning could be a function of different task / behavioral demands. The delayed-win-shift radial arm maze task requires greater spatial memory than does shuttling for food rewards (e.g., circle track running). Further, in comparison to track-running testing, rats receive unusually extensive familiarization with the radial arm maze, albeit with a random set of arms open during the study phase, as they complete daily testing for two to four months during shaping.

The purpose of the current study was to investigate the sufficiency of wall height for explaining differences in the expression of head scanning and rearing behaviors. To test this hypothesis, we fixed the behavioral context to be the delayed win-shift radial arm maze task but varied the wall height from one trial to the next. We hypothesize that, if head scanning and rearing are functionally related behaviors, then rats will transition from rearing to head scanning when the height of the walls is dropped. Moreover, if the rats treat the visually distinct behaviors as interchangeable, we would expect that the specifics of where and when the rats head scan will align the rearing behavior when the walls were tall. Better understanding of these behaviors will serve to guide future studies examining unique epochs of hippocampal function.

## Methods

All procedures and surgeries were conducted in strict accordance with National Institutes of Health and under a protocol approved by the Indiana University Bloomington Institutional Animal Care and Use Committee (IU BIACUC).

### Subjects

A total of 9 adult Long Evans rats were used in this study (4 males, 5 females). Animals were pair-housed, given ad libitum access to water, maintained at 85%-90% of free-feeding weight, and maintained on a 12-h light/dark cycle. Before testing, rats were trained to criterion on the delayed spatial win-shift radial arm maze task using the same apparatus and procedure as we have used previously repeated here for transparency (Layfield et al., 2023; Cassity et al., 2024).

### Apparatus

Behavioral training took place in a custom 8-arm radial maze with computer-controlled pneumatic drop doors at the entrances. The maze had a 33.2cm-wide hub and each arm measured 48.26 cm-long, 10.79 cm-wide with 20.95cm-tall walls. Walls were made from clear acrylic to allow for visual orientation to distal cues that surrounded the maze. Walls in short-wall sessions were 4.6 cm tall and made of opaque matte white acrylic. The floors were made from opaque matte white acrylic. At the end of each arm were food wells in which 45-mg chocolate flavored sucrose pellets (Bio-Serv, Flemington, NJ) were delivered. The maze was situated in a 3 m by 3.6 m room, surrounded by rich visual cues to facilitate allocentric orientation. Cues were constructed from brightly colored tape or construction paper with a row of six cues evenly spaced level with the walls of the maze and a second row of cues approximately 2 ft above the first row of cues. The room also contains a framed image, a desk, a bookshelf, and the holding pedestal on which rats were placed between the trial phases immediately adjacent to the maze (see photo in Figure 1). A high-definition camera (FLIR Flea3 or Intel RealSense D435) mounted to the ceiling above the maze was connected to a PC running Open Broadcaster Software (OBS) to record the movements of the rat for offline analysis.

**Figure 1:**
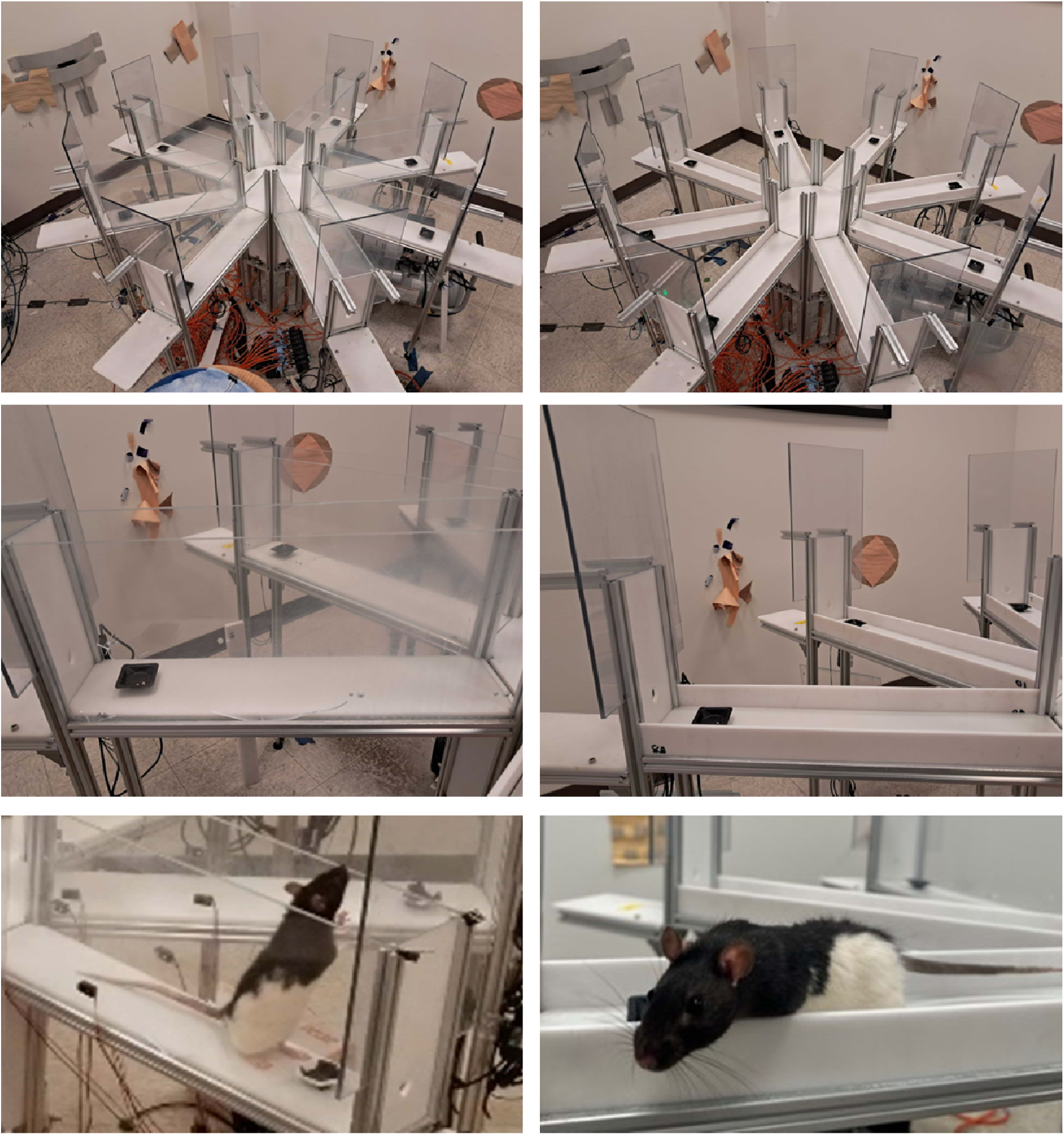
Left, images of the eight-arm radial maze with tall walls. Bottom image shows an example of a rearing event. Right, images of the eight-arm radial maze with short walls. Bottom image shows an example of a head scanning event.

### Behavioral Training

#### Habituation and preliminary training

Before any behavioral training, all rats were handled daily for 3-5 minutes for one week. Animals then underwent a habituation phase during which they were placed in the maze for 15 minutes daily for five days. During these sessions, a small weigh boat at the end of each arm was baited with two 45 mg sucrose pellets. Pellets were also spread evenly throughout the floor of the maze to encourage exploration. After each session, maze floors, walls, and doors were cleaned with 10% chlorhexidine solution to ensure cleanliness of the maze and reduce potential odors cues.

After the habituation phase, animals then moved on to the preliminary training phase. In this phase, animals were placed in the maze hub for 30 seconds, after which all doors opened. All arms were baited with two sucrose pellets and the session lasted for either 15 minutes or until each arm was entered and all pellets were consumed. During this time, all arm entries were recorded, with an entry being defined as the rump of the animal entering the arm. Animals completed this training daily until reaching a criteria of, over four consecutive days, no more than two errors each day. Errors were defined as re-entry to an arm already visited that day.

#### Delayed win-shift task

After reaching criteria in the preliminary training phase, animals then moved on to the final delayed win-shift task. Before the animal was brought into the room, all maze arms and doors were cleaned with chlorhexidine and the small weigh boat at the end of each arm was baited with two sucrose pellets. The animal was then brought into the room and placed on a pedestal next to the maze. The animal was then placed in the middle hub of the maze with all doors closed and waited for 30 seconds. The study phase of the task then began, during which the doors to four random arms (changing daily) opened. The animal then freely navigated the maze until all pellets were retrieved from the four open arms. After which they were taken out of the maze and placed back on the pedestal outside of the maze for a delay of 1-2 minutes. During the delay the maze was again cleaned with chlorhexidine. The animal was then again placed in the middle hub of the maze with all doors closed for 30 seconds. At this point, the test phase began, during which all doors opened and the animal was tasked with navigating to the four arms that it did not travel to in the study phase. That is, the animal was challenged to retain in its working memory the four arms from the study phase, so that it could efficiently forage and enter the other four arms in the test phase. After retrieving the pellets in the remaining arms, the animal was removed from the maze and the training session ended. Regular training (∼5 days/week) continued until rats achieved the behavioral criterion. Criterion was defined as no more than two errors in the test phase and no errors in the study phase over four days. Errors were defined as re-entry to an arm already visited that day.

### Behavioral Testing and Experimental Conditions

Once animals reached criteria on the task, animals completed six short-wall habituation days. Short-wall habituation involved running the rat per the same protocol as the standard delayed win-shift task, one trial per day, but the standard 21 cm tall walls along the sides of each arm were replaced with 4.6 cm short walls during the study phase only. Based on the expectation that the novelty of these walls would affect the behavior, rearing and head scanning was not examined on these days. Key data collection was performed on the subsequent 16 days, treated as eight pairs of two days. Within each pair, the order of short versus tall walls was randomized. This pattern allowed for eight days of data collection in each wall condition. As in the habituation phase, short walls were only ever used during the study phase

### Behavioral Scoring

Videos were recorded using Open Broadcaster Software (OBS) Studio. Animal behavior was manually scored using VirtualDub2 and DJV to find the frame numbers in which key behaviors (rearing and head scanning) started and ended. Rearing initiation was defined as the moment the forelegs of the rat left the ground. Rearing termination was the moment the forelegs returned to the maze floor. Head scanning started when the head of the rat broke the plane of the wall and ended when the head ceased to break the plane of the wall. Analyses that were standardized by time used the length of the corresponding phase: Data from the study phase were standardized by the study phase duration and test phase data were adjusted by the test phase duration.

### Analysis

#### Animal tracking

Analyses of the rat position at the time of exploratory behaviors leveraged tracking provided by applying DeepLabCut (Mathis et al., 2018; Nath et al., 2019) to video recordings of the rat in the maze collected from the overhead camera (FLIR Flea3 or RGB sensor of an Intel RealSense D435). A DeepLabCut network was built with 24 frames from each of 10 rats (240 total frames) from a prior cohort labelled to mark the nose, left ear, right ear, middle of the back, and tail base (rump). After an initial 300-k training iterations, an additional 240 ‘jumpy’ frames were extracted, relabeled, and added to the training set. The network then received another 600-k training iterations on the expanded training set. Network performance was then confirmed by manual inspection by checking labelled videos and quantifying jumpy frames in the extracted tracking data. Prior to use, the tracking data were preprocessed to interpolate over jumps of >5 cm from one frame to the next. Rat position at the time of key behaviors was tracked the position of the middle of the back. This was used to better estimate the central position of the rat, as opposed to head position which may vary a lot especially during head scanning.

#### Behavior initiation rate maps

The spatial distributions of key behaviors were obtained by aligning the start and end frame numbers to the animal tracking records. Behavior initiation rate maps were created using the standard procedures for creating neural firing rate maps: Binned behavior initiation count maps (with 1.8 cm bins) were smoothed and then standardized by smoothed occupancy maps. The smoothing was performed with a Gaussian kernel with sigma set to 2 bins (3.6 cm).

#### Bootstrapping

The spatial distributions where key behaviors (rearing or head scanning) were initiated were compared to null distributions generated through bootstrapping methods. The bootstrap samples were generated by applying a pseudo-random temporal offset to the onset times of the respective key behavior to select new random moments when the rat could have initiated the behavior. Frames where the animal was in the key behavior were removed from the set of frames treated as times of potential key behavior initiation. The temporal offsets were constrained to be 20-80% of the trial duration. The null distribution was calculated separately for each trial from 500 iterations. From each, the mean and standard deviation over spatial bins was saved and used to z-score the empirically observed behavior initiation likelihoods. This method randomizes the relationship between the onset of key behaviors and the position of the rat. Behavioral distance distributions were smoothed across spatial bins using a moving average filter applied to the binned values, reducing high-frequency noise while preserving broader spatial trends. Smoothing was performed within each animal prior to aggregation to ensure that local variability did not disproportionately influence group-level comparisons.

#### Statistical Testing

To quantify differences in behavioral measures across conditions (Figure 3), nonparametric statistical tests were used due to a lack of normality. Friedman tests were used as an omnibus method for initially detecting significant differences between conditions. When a significant effect was detected using the Friedman test, post hoc pairwise comparisons were performed using Wilcoxon signed-rank tests. These tests assessed differences between conditions in a pairwise manner. To control for multiple comparisons, p-values were adjusted using Bonferroni correction based on a significance level of α = 0.05.

**Figure 2:**
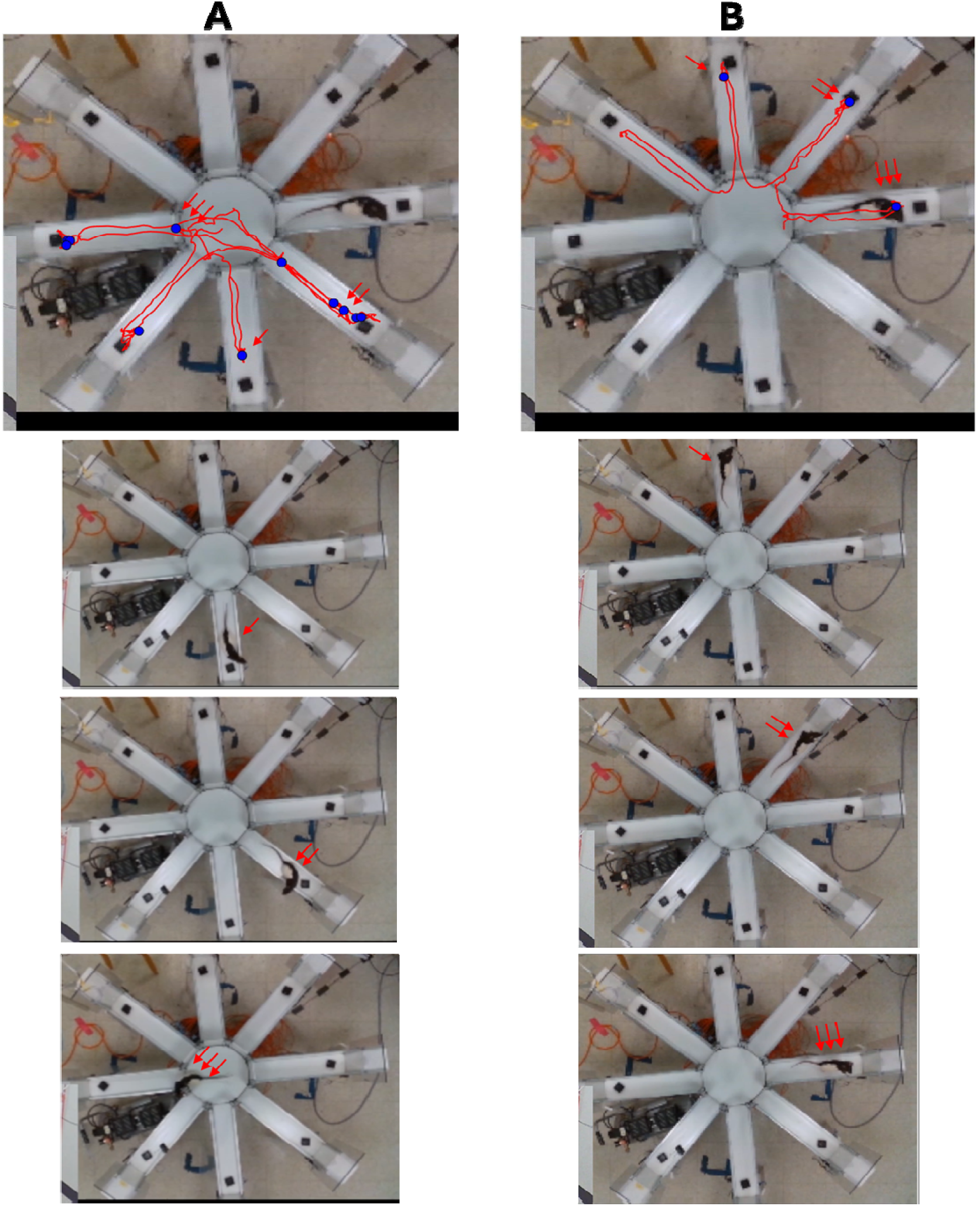
Example behaviors and animal trajectory. A) Top – For one testing session, the animal trajectory (red line) and head scanning events (blue circles) during the study phase. Bottom – Images of actual head scanning events. Red arrows correspond to the locations in the top image. B) Top – For one testing session, the animal trajectory (red line) and rearing events (blue circles) during the test phase. Bottom – Images of actual rearing events. Red

**Figure 3:**
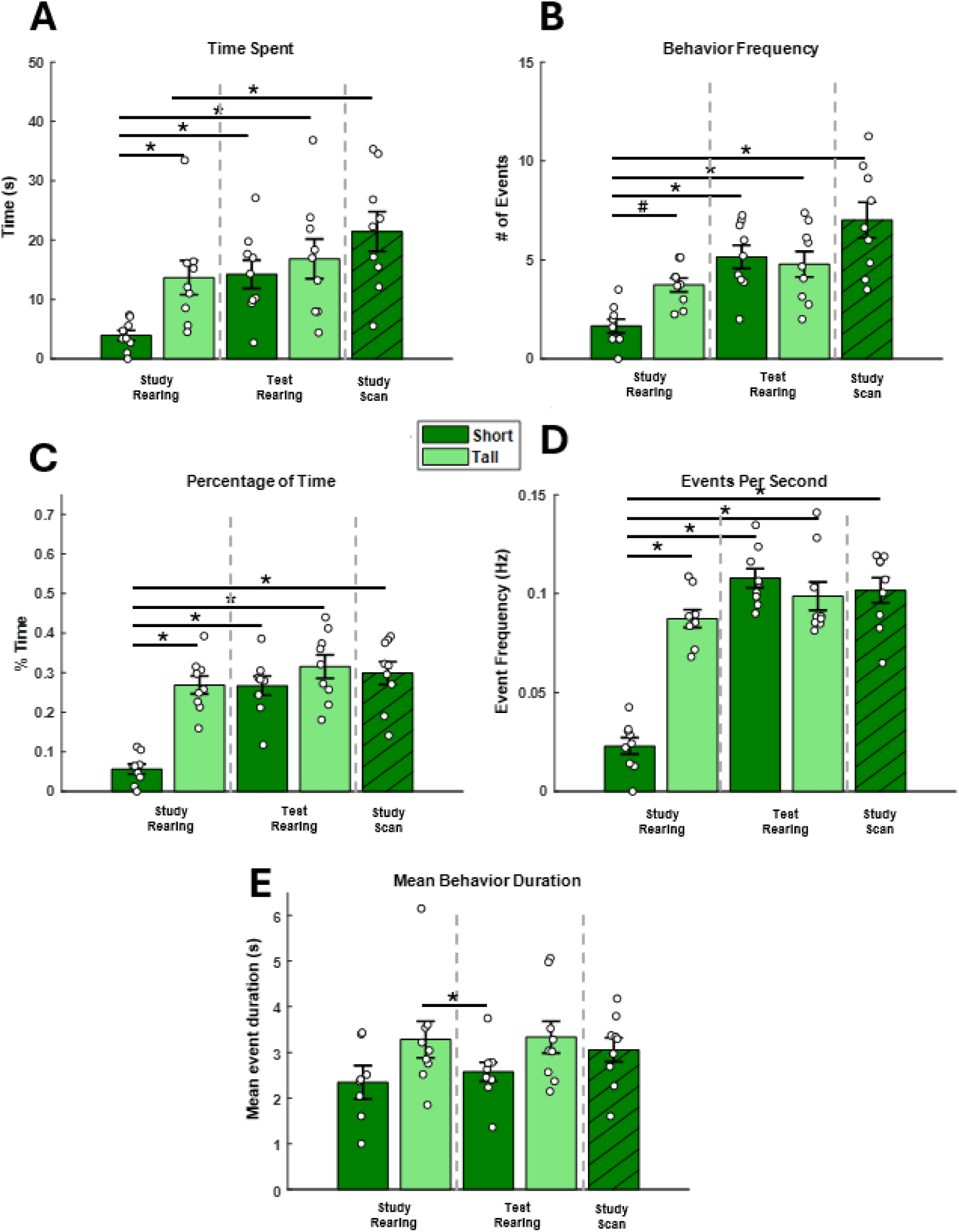
Behavioral metrics comparing rearing and scanning according to experimental phase and wall type. A) Mean total time spent rearing or LHS. B) Mean number of rearing and scanning events. C) Mean percentage of phase time spent in each behavior. D) Mean frequency adjusted by phase length, shown as event frequency. E) Mean behavior duration (i.e. total behavior time divided by total behavior events). Note for all A-E, the x axis shows the conditions being quantified and bar color indicates wall type. For example, “Study Rearing” includes rearing data from the study phase and bar color indicates the wall type for those sessions. Error bars indicate the standard error of the mean and white circles surrounding each bar are data points for individual animals (A-E: n = 9). (* indicates p < 0.05; # indicates p < 0.1)

To assess whether the overall spatial distribution of behavior differed from the null distribution (Figure 4B), we computed the Wasserstein distance between the mean true and mean null distributions across distance bins. Statistical significance was evaluated using a permutation test in which, for each iteration, true and null labels were randomly swapped within each animal and the Wasserstein distance recomputed to generate a null distribution of distances. The observed Wasserstein distance was then compared to this distribution to obtain a p-value, providing an omnibus test of whether the true spatial distribution differs from that expected under the null.

**Figure 4:**
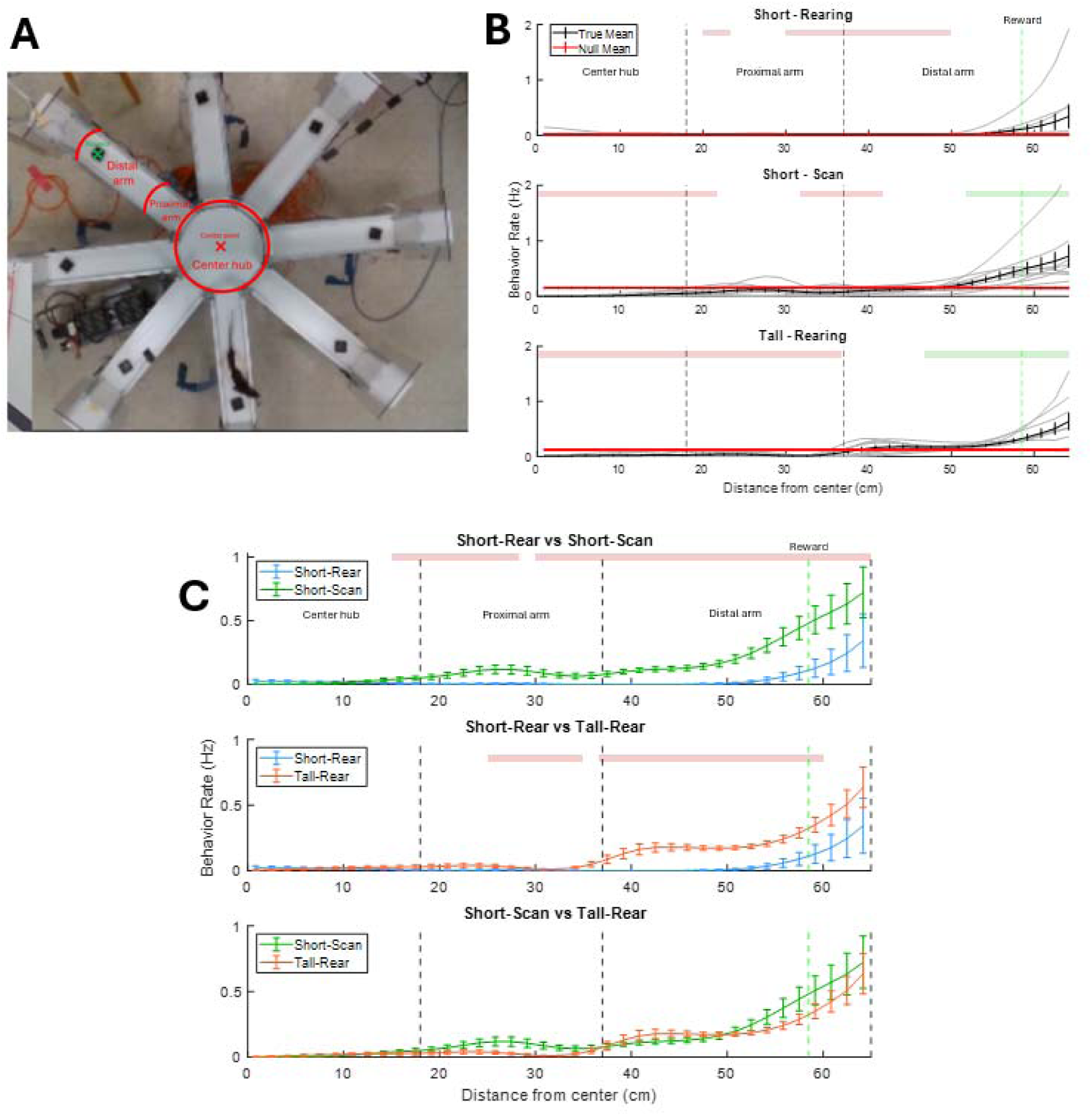
Spatial distribution of rearing and scanning measured as behavioral rate. A) Illustration showing location and terminology used to demarcate maze areas used in analysis. Red lines indicate borders between maze zones and green “x” indicates reward site. B) Comparison of observed behavioral spatial distributions (black line) compared to a null distribution computed by bootstrapping (red line). C) Pairwise comparison of behavioral conditions. For both B and C, red/green shaded bars at the top of each graph indicate significant differences between the two distributions at those bins (p < 0.05). Red indicates the condition listed at the top of the legend is significantly lower compared to the condition below it, while green indicates the opposite. Black dashed lines indicate borders of maze zones. Green dashed line indicates reward site. Error bars indicate standard error of the mean across rats for each bin.

To test for overall differences between conditions across spatial bins (Figure 4C), we used a Friedman test. For each animal, values from the two conditions were treated as repeated measures across spatial bins, and the Friedman test assessed whether there was a consistent difference in distributions across bins at the group level. This approach does not assume normality and accounts for the repeated-measures structure of the data.

To identify spatial regions in which behavioral distributions differed either from a null distribution (Figure 4B) or between experimental conditions (Figure 4C), z-scores between the values were first calculated on a per-trial basis for each bin. Per-rat maps were created for each condition by averaging z-scores over trials. The mean and standard error of the mean of these z-scores were then calculated across rats and a one-sample t -test was performed for each bin. A Benjamini-Hochberg False Discovery Rate (FDR) correction was then applied to adjust for multiple comparisons and α = 0.05 was used to determine significance.

## Results

### Decreased wall height leads to shift from rearing to head scanning

To understand how wall height affects rearing and head scanning behavior, we split the data by phase type (study and test), wall type (short and tall), and behavior type (rear and head scan). Note that wall height was only changed in the study phase of the experiment. However, data from the test phase was included as a comparison to ensure behavior was unchanged under normal wall conditions following exposure to short walls in the same session.

First, the mean total time spent rearing and head scanning was examined. A Friedman test revealed a significant difference in time spent rearing and head scanning across conditions (χ²_4_ = 23.82, p < 0.001). Post-hoc Wilcoxon signed-rank tests with Bonferroni correction indicated that time spent rearing in the study phase of short-wall sessions (3.93 ± 0.85) was significantly lower than all other conditions (all p < 0.05). (Figure 3A).

Next, we assessed the mean number of rearing and head scanning events across conditions. A Friedman test revealed a significant difference in the number events across conditions (χ²_4_ = 25.18, p < 0.001). Post-hoc Wilcoxon signed-rank tests with Bonferroni correction indicated that the number of rearing events in the study phase of short-wall sessions (1.66 ± 0.35) was significantly lower compared to all other conditions (p < 0.05) with the exception of the tall-wall study phase, for which the difference was a trending decrease (p = 0.078) (Figure 3B).

To ensure these effects were not the result of differences in the duration of trial phases, we analyzed the percentage of time spent rearing or head scanning relative to the phase length. The results were qualitatively the same. A Friedman test revealed a significant difference in the number events across conditions (χ²_4_ = 22.49, p < 0.001). Post-hoc Wilcoxon signed-rank tests with Bonferroni correction indicated that percent time spent rearing was significantly decreased in the study phase of short-wall sessions (0.06 ± 0.01) compared to all other conditions (all p < 0.05) (Figure 3C).

We then adjusted the number of behavioral events by phase length and found similar results. A Friedman test revealed a significant difference in the number events across conditions (χ²_4_ = 22.49, p < 0.001). Post-hoc Wilcoxon signed-rank tests with Bonferroni correction indicated that rearing was significantly decreased in the study phase of short-wall sessions (0.02 ± 0.004) compared to all other conditions (all p < 0.05) (Figure 3D).

Lastly, we examined the mean duration of individual behavioral events to see if wall or phase type affected how long animals performed each behavior once initiated. A Friedman test revealed a significant difference in the number events across conditions (χ²_4_ = 12.5, p < 0.05). Post-hoc Wilcoxon signed-rank tests with Bonferroni correction indicated that the mean duration of individual rears was significantly shorter in the test phase of short-wall sessions compared to that seen in the study phase of tall-wall sessions (p < 0.05) (Figure 3E).

Taken together, these results indicate that rats rear less when environmental boundaries are shorter. Importantly, however, rearing is not completely abolished by the change to short walls. In addition, head scanning behavior was comparable to rearing behavior in tall-wall sessions (all comparisons p > 0.1), suggesting head scanning largely replaces rearing behavior when short walls are present.

### Wall type affects spatial distribution of exploratory behaviors

To further explore the possibility that rats treat rearing and head scanning as interchangeable, replacing rearing with head scanning when permitted, we examined and compared where rats performed these behaviors under each circumstance.

To achieve this, we analyzed the spatial distribution of where each behavior occurred on a linearized position, calculated as the rate of behavior at each bin. The linearized position was established as the distance from the center of the maze. We then used a bootstrapping method in which, over 500 iterations, the onset timestamp of each behavior was shifted randomly. This provided a simulated null distribution to which the observed data was compared. We then computed z-scores for each rat between the observed and null values at each bin in each condition and these z-scores were then averaged across rats. To determine significance, a one-sample t-test was then run for each bin and a Benjamini-Hochberg False Discovery Rate (FDR) correction was used to correct for multiple comparisons.

For short-wall rearing, the observed distribution differed significantly from the null distribution (Wasserstein distance = 20.15, *p* = 0.122). Although the global distribution was not significantly different from the null, 14 of 39 bins were significant, forming two distinct spatial segments (20–23.33 cm and 30–50 cm), both reflecting reduced rearing relative to the null (*p* = 0.0078 and 0.0013, respectively). These findings indicate localized suppression of rearing during short-wall sessions despite the absence of a global shift (Figure 4B).

For head scanning in short-wall sessions, the observed global distribution did significantly differ from the null (Wasserstein distance = 17.84, *p* = 0.006). In addition, 27 of 39 bins were significantly different from the null, forming three spatial segments. Two segments (0.00–21.67 cm and 31.67–41.67 cm) showed reduced scanning relative to the null (*p* = 0.0026 and 0.0132, respectively), while a distal segment (51.67–65.00 cm) showed increased scanning behavior (mean *p* = 0.0016) (Figure 4B).

For tall-wall rearing, the observed distribution again differed significantly from the null (Wasserstein distance = 18.49, *p* = 0.004). Further examination of localized differences showed that 33 of 39 bins were significantly different from the null, forming two broad spatial segments. A proximal segment (0.00–36.67 cm) exhibited reduced rearing relative to the null (p = 0.0002), whereas a distal segment (46.67–65.00 cm) showed increased rearing (*p* = 0.0051) (Figure 4B).

There are a few notable takeaways from these results. First, rearing in short-wall trials was significantly reduced throughout a large portion of the proximal and distal maze arms. Second, head scanning in the short-wall trials was biased towards the end of the distal arm area and was significantly lower in the center hub and beginning portions of the maze arms. Lastly, rearing frequency in the tall-wall condition was significantly lower in the center hub and proximal arm areas, while it was significantly higher in the distal arm areas. Together, these results indicate that wall height has a significant effect on the magnitude and the spatial distribution of rearing and head scanning behavior. In particular, rearing and head scanning occur in a biased fashion around reward sites. However, the rearing bias was only present in tall-wall sessions and this behavior seemed to be replaced by head scanning in short-wall sessions.

Next, we performed the same analysis, but instead directly compared the spatial distributions of each behavior across conditions. Friedman tests were used to first assess overall differences between each distribution. Localized differences were then examined using the same z-scoring method as previously described.

Comparing short-wall rearing and short-wall head scanning, there was a significant overall difference across spatial bins (Friedman χ²(1) = 9.00, *p* = 0.0027). A large proportion of bins were significant after FDR correction (29/39), forming two contiguous spatial segments. Specifically, scanning behavior exceeded rearing behavior across both an intermediate region (15.00–28.33 cm; *p* = 0.0356) and a broad distal region (30.00–65.00 cm; *p* < 0.001), indicating a robust shift toward increased scanning relative to rearing across much of the spatial domain (Figure 4C).

A similar pattern emerged when comparing short-wall rearing to tall-wall rearing. There was a significant overall difference between conditions (Friedman χ²(1) = 9.00, *p* = 0.0027), with 20 of 39 bins reaching significance following FDR correction. These significant bins formed two spatial segments, including a region in the proximal arm (25–35 cm; *p* = 0.0405) and a larger distal region (36.67–60 cm; *p* = 0.0015). In both regions, tall-wall rearing exceeded short-wall rearing, suggesting a spatially selective increase in rearing behavior under the tall condition (Figure 4C).

In contrast, comparison of short scanning and tall rearing revealed no significant differences. The Friedman test did not detect an overall effect (χ²(1) = 1.00, *p* = 0.317), and no bins survived FDR correction (0/39). Accordingly, no spatially contiguous segments of significant difference were observed, indicating that these two conditions produced comparable spatial distributions of behavior (Figure 4C).

Together, these findings demonstrate that spatial behavioral profiles differ systematically according to wall type. In particular, rearing under short-wall conditions is significantly reduced, including in the distal arm area where rearing is most robust during tall-wall conditions. Head scanning in the short-wall condition also seemed to replace rearing at this distal arm location, suggesting that rats switch to head scanning when short walls are present.

## Discussion

The current study examined how rat exploratory behavior changes in response to boundary height. The rearing and head scanning behaviors of rats proficient in the radial arm maze delayed win-shift task were examined when the height of the maze walls was manipulated. We hypothesized that, when the standard tall walls were replaced with short walls, the rats would head scan instead of rear but that the specifics of the head scanning would match those of the rearing when the walls had been tall. Both hypotheses were supported by our observations, suggesting that the boundary height is a significant factor in determining which exploratory behavior the rats express. These results indicate that rats may treat rearing and head scanning as interchangeable.

This experiment aimed to fill a key gap in what is known regarding the relationship between these two exploratory behaviors. Rearing is a period of hippocampal dependent spatial memory encoding in a radial arm maze delayed win-shift task (Layfield et al., 2023) and head scanning is associated with the creation of new place fields in hippocampal neurons (Monaco et al., 2014). Both exploratory behaviors reflect attentive information gathering and are generated in response to environmental novelty, raising the possibility that they are posturally distinct variations of single functional behavior. The implication of which is that rearing would also be a time of place field formation and head scanning would be an epoch of hippocampal dependent spatial memory encoding. The results of this experiment give reason to believe that rearing and head scanning are indeed functionally similar.

If rearing marks points of likely place field formation, as does head scanning, it would be significant. The mechanisms of hippocampal spatial coding remain a highly active area of research with seemingly basic questions remaining poorly addressed. This includes questions about why place fields are distributed as they are and regarding the conditions under which spatial tuning evolves as a function of experience. Here, we showed that rearing and head scanning occurs disproportionately often near reward sites at arm ends, a pattern that could explain the clustering of place fields near reward sites (Hollup et al., 2001; Dupret et al., 2010; Gauthier & Tank, 2018; Jarzebowski et al., 2022). This informs our understanding of the behavioral conditions that drive the evolution of hippocampal coding. However, it remains ambiguous from this work whether the exploration was tied to the rewards sites or the arm ends. While the reward sites have direct task relevance, the arm ends are areas of higher availability of sensory information discriminative of allocentric position. The latter explanation aligns with the findings of Davis et al. (2025) who found head scanning broadly distributed across a circular track in a room rich in visual landmarks instead of clustered near the reward sites. Further work is required to A) verify whether place tuning is updated during rearing, and B) to better understand the behavioral contexts that motivate attentive sampling.

If head scanning is an epoch of hippocampal dependent spatial memory encoding, as rearing is, it would also be significant. Despite the substantial attention dedicated to understanding hippocampal function, the set of contexts in which key hippocampal dependent learning occur during is short. Moments of locomotion and sharp wave ripples have dominated the consensus view of the key epochs of hippocampal function (O’Keefe & Dostrovsky, 1971; Knierim et al., 1995; Ekstrom et al. 2003; Buzsaki et al. 2013; Diba & Buzsaki, 2007; Jadhav et al., 2012; Csicsvari & Dupret D, 2014; Gillespie et al. 2021; cf. Monaco et al., 2014). Yet, if head scanning, like rearing, supports hippocampal dependent memory, it would indicate that attentive sampling, more broadly, is a third key epoch of hippocampal processing. Such behaviors have received disproportionately little attention in hippocampal research in recent years and, indeed, are often proactively filtered from recorded datasets during analyses of hippocampal function. The current work, together with other evidence, supports turning renewed focus on the importance of these attentive sampling behaviors.

Both rearing and head scanning are associated with elevated hippocampal theta power (Vinogradova 1995). The current source density profile of theta during these behaviors is, however, distinct from that observed during running in the case of rearing at least. Barth et al. (2018) revealed theta during rearing to be a period during which input from the medial entorhinal cortex layer II to the dentate gyrus of the hippocampus is strengthened (Barth et al., 2018). This phasic increase suggests that the hippocampus enters a functionally distinct state during rearing. Such an assessment of hippocampal dynamics has not been performed for epochs of attentive head scanning. The increased entorhinal input to the dentate gyrus during rearing is consistent with the view of rearing, and putatively head scanning, as epochs of overt sensory sampling.

The current results should be considered in the context of what type of sensory information is being collected. When rats rear in tall-wall sessions, one possibility is that they are attempting to access olfactory airstreams above the maze walls. In most cases rats orient themselves vertically during rearing in these conditions, with the nose pointed upwards; although, to our knowledge sniffing has not been quantified in this context. However, some studies do suggest rats may target olfactory stimuli during rearing. For instance, Dielenberg & McGregor (2001) use a behavioral paradigm where they show that rats rear in an oriented fashion towards a cat odor source. In addition, rats may be using their own smell from recent visits to other parts of the maze to navigate, as Wallace et al. (2002) shows that rats can distinguish their own odor from other rat odors. The current results may suggest that during short-wall sessions, rats were able to access olfactory information without rearing and therefore opted for head scanning. A useful future study would be to quantify sniffing during head scanning and rearing in the current experiment.

Another possibility is that during rearing, rats are attempting to look over the wall to investigate visual stimuli in the room. While clear Plexiglas walls are used in our experiment, it is likely they still obscure visual information to an extent. Animals may therefore, similar to olfactory information, attempt to better access visual information by rearing. Previous work does suggest that rats prefer to use visual information during navigation when it is available (Lavenex & Schenk, 1995; Lavenex & Schenk, 1997; Maaswinkel & Whishaw, 1999). In addition, there is a wealth of literature showing how spatially-tuned cells respond to changes in contextual visual stimuli (e.g. O’Keefe & Conway, 1978; Muller & Kubie, 1987; Fenton et al., 2000a,b; Scaplen et al., 2014). Future work should investigate what sensory information is being targeted during rearing.

Importantly, however, most animals did not completely abandon rearing during short wall trials. This indicates that, although head scanning may replace most rearing behavior during short wall trials, rearing was still used in some rare cases. An important note here is that, in the short wall condition, the wall at the end of each arm remained tall. This may account for the small amount of rearing still occurring in the short-wall sessions and may explain why rearing in this condition appears slightly shifted towards the end of the arm.

The observation that rearing and head scanning may not be fully interchangeable to the rats bears relevance for inferences about the upstream physiological mechanisms that motivate these behaviors. These mechanisms are poorly understood for both behaviors. We previously found that, despite strong correlations in freely behaving rats between population activity of cholinergic neurons in the medial septum and diagonal band of Broca and rearing (Kopsick et al., 2022), that optogenetic inhibition of these neurons does not reduce rearing frequency (Cassity, Choi, et al., 2024). The inhibition did, however, result in individual rears being longer as though it slowed encoding of the information being sampled. What has been shown to motivate rearing is the activity of melanin-concentrating hormone neurons of the lateral hypothalamus (Concetti et al., 2024) and glutamatergic neurons of the medial septum that target the ventral tegmental area (Mocellin et al., 2024). Considerably more work is required to understand when and how these circuits modulate environmental sampling and exploration.

A limitation of the current study is that all rats had been trained to proficiency on the maze with tall walls prior to the current testing. Thus, the current results do address whether the head scanning behavior may be different in rats trained with short walls from the outset. Nonetheless, it remains evident from the current results that rats preferentially head scan when the walls are short but that most rats will not abandon rearing completely. A second limitation of the current study is that it did not manipulate task demands. Rats were subject to similar task demands in both conditions, those of the delayed spatial win shift task, and all were highly familiarized with the radial arm maze prior to testing. Yet, the current data nonetheless demonstrate sufficiency of manipulating wall height for affecting the type of exploratory behaviors rats expressed.

In summary, the current study examined how wall height influenced the frequency of rearing and head scanning. Taken together, results suggest that rearing and head scanning may have similar underlying hippocampal functions and occur depending on environmental layout. These behaviors occur predominantly around the reward sites, suggesting involvement in hippocampal spatial processing. This study sets the stage for further experiments to investigate hippocampal function during rearing and head scanning to further understand their similarities and differences.

## References

Barth, A. M., Domonkos, A., Fernandez-Ruiz, A., Freund, T. F., & Varga, V. (2018). Hippocampal Network Dynamics during Rearing Episodes. Cell Reports, 23(6), 1706–1715. 10.1016/j.celrep.2018.04.021

Boccara, C. N., Sargolini, F., Thoresen, V. H., Solstad, T., Witter, M. P., Moser, E. I., & Moser, M.-B. (2010). Grid cells in pre- and parasubiculum. Nature Neuroscience, 13(8), 987–994. 10.1038/nn.2602

Buzsáki, G. & Moser, E. I. Memory, navigation and theta rhythm in the hippocampal-entorhinal system. Nature Neuroscience 16, 130–138 (2013).

Cassity, S., Choi, I. J., Gregory, B. H., Igbasanmi, A. M., Bickford, S. C., Moore, K. T., Seraiah, A. E., Layfield, D., & Newman, E. L. (n.d.). Cholinergic modulation of rearing in rats performing a spatial memory task. European Journal of Neuroscience, n/a(n/a). 10.1111/ejn.16248

Concetti, C., Viskaitis, P., Grujic, N., Duss, S. N., Privitera, M., Bohacek, J., Peleg-Raibstein, D., & Burdakov, D. (2024). Exploratory Rearing Is Governed by Hypothalamic Melanin-Concentrating Hormone Neurons According to Locus Ceruleus. The Journal of Neuroscience, 44(21), e0015242024. 10.1523/jneurosci.0015-24.2024

Csicsvari, J., & Dupret, D. (2014). Sharp wave/ripple network oscillations and learning-associated hippocampal maps. Philosophical Transactions of the Royal Society B: Biological Sciences, 369(1635), 20120528. 10.1098/rstb.2012.0528

Davis, P. J., Jones, S. T., & Savelli, F. (2026). Prevalence and modulation of rat off-track head-scanning on linear tracks: possible implications for representational and dynamical properties of hippocampal place cells. bioRxiv, 2025.05.27.656456. 10.1101/2025.05.27.656456

Diba K, Buzsáki G (2007) Forward and reverse hippocampal place-cell sequences during ripples. Nature Neuroscience, 10(10):1241–1242. 10.1038/nn1961

Dielenberg, R. A., & McGregor, I. S. (2001). Defensive behavior in rats towards predatory odors: A review. Neuroscience & Biobehavioral Reviews, 25(7), 597–609. 10.1016/S0149-7634(01)00044-6

Dupret, D., O’Neill, J., Pleydell-Bouverie, B. & Csicsvari, J. The reorganization and reactivation of hippocampal maps predict spatial memory performance. Nature Neuroscience 13, 995–1002 (2010).

Ekstrom AD, Kahana MJ, Caplan JB, Fields TA, Isham EA, Newman EL, Fried I (2003) Cellular networks underlying human spatial navigation. Nature, 425(6954):184–188. 10.1038/nature01964

Fenton, A. A., Csizmadia, G., & Muller, R. U. (2000a). Conjoint Control of Hippocampal Place Cell Firing by Two Visual Stimuli: I. the Effects of Moving the Stimuli on Firing Field Positions. Journal of General Physiology, 116(2), 191–210. 10.1085/jgp.116.2.191

Fenton, A. A., Csizmadia, G., & Muller, R. U. (2000b). Conjoint Control of Hippocampal Place Cell Firing by Two Visual Stimuli: II. a Vector-Field Theory That Predicts Modifications of the Representation of the Environment. Journal of General Physiology, 116(2), 211–222. 10.1085/jgp.116.2.211

Ferbinteanu, J., Ray, C., & McDonald, R. J. (2003). Both dorsal and ventral hippocampus contribute to spatial learning in Long–Evans rats. Neuroscience Letters, 345(2), 131–135. 10.1016/S0304-3940(03)00473-7

Gauthier, J. L., & Tank, D. W. (2018). A Dedicated Population for Reward Coding in the Hippocampus. Neuron, 99(1), 179–193.e7. 10.1016/j.neuron.2018.06.008

Gillespie, A. K., Maya, D. A. A., Denovellis, E. L., Liu, D. F., Kastner, D. B., Coulter, M. E., Roumis, D. K., Eden, U. T., & Frank, L. M. (2021). Hippocampal replay reflects specific past experiences rather than a plan for subsequent choice. Neuron, 109(19), 3149–3163.e6. 10.1016/j.neuron.2021.07.029

Hafting, T., Fyhn, M., Molden, S., Moser, M.-B., & Moser, E. I. (2005). Microstructure of a spatial map in the entorhinal cortex. Nature, 436(7052), 801–806. 10.1038/nature03721

Hollup, S. A., Molden, S., Donnett, J. G., Moser, M.-B. & Moser, E. I. (2001) Accumulation of Hippocampal Place Fields at the Goal Location in an Annular Watermaze Task. J. Neurosci. 21, 1635–1644.

Jadhav SP, Kemere C, German PW, Frank LM (2012) Awake hippocampal sharp-wave ripples support spatial memory. Science (New York, NY), 336(6087):1454–1458. 10.1126/science.1217230

Knierim, J., Kudrimoti, H. & McNaughton, B. Place cells, head direction cells, and the learning of landmark stability. J Neurosci 15, 1648–1659 (1995).

Kopsick, J. D., Hartzell, K., Lararo, H., Nambiar, P., Hasselmo, M.E., Dannenberg, H. (2022) Temporal dynamics of cholinergic activity in the septo-hippocampal system. Front Neural Circuit 16, 957441. 10.3389/fncir.2022.957441

Lavenex, P., & Schenk, F. (1995). Influence of local environmental olfactory cues on place learning in rats. Physiology & Behavior, 58(6), 1059–1066. 10.1016/0031-9384(95)02002-0

Lavenex, P., & Schenk, F. (1997). Olfactory Cues Potentiate Learning of Distant Visuospatial Information. Neurobiology of Learning and Memory, 68(2), 140–153. 10.1006/nlme.1997.3791

Layfield, D., Sidell, N., Blankenberger, K., & Newman, E. L. (2023). Hippocampal inactivation during rearing on hind legs impairs spatial memory. Scientific Reports, 13(1), 6136. 10.1038/s41598-023-33209-9

Lever, C., Burton, S., Jeewajee, A., O’Keefe, J., & Burgess, N. (2009). Boundary Vector Cells in the Subiculum of the Hippocampal Formation. Journal of Neuroscience, 29(31), 9771–9777. 10.1523/JNEUROSCI.1319-09.2009

Lever, C., Burton, S., & Ο’Keefe, J. (2006). Rearing on Hind Legs, Environmental Novelty, and the Hippocampal Formation. Reviews in the Neurosciences, 17(1–2), 111–134. 10.1515/revneuro.2006.17.1-2.111

Maaswinkel, H., & Whishaw, I. Q. (1999). Homing with locale, taxon, and dead reckoning strategies by foraging rats: Sensory hierarchy in spatial navigation. Behavioural Brain Research, 99(2), 143–152. 10.1016/S0166-4328(98)00100-4

Mathis, A., Mamidanna, P., Cury, K. M., Abe, T., Murthy, V. N., Mathis, M. W., & Bethge, M. (2018). DeepLabCut: Markerless pose estimation of user-defined body parts with deep learning. Nature Neuroscience, 21(9), 1281–1289. 10.1038/s41593-018-0209-y

Mocellin P, Barnstedt O, Luxem K, Kaneko H, Vieweg S, Henschke JU, Dalügge D, Fuhrmann F, Karpova A, Pakan JMP, Kreutz MR, Mikulovic S, Remy S. (2024). A septal-ventral tegmental area circuit drives exploratory behavior. Neuron 112:1020–1032.e7.

Monaco, J. D., Rao, G., Roth, E. D., & Knierim, J. J. (2014). Attentive scanning behavior drives one-trial potentiation of hippocampal place fields. Nature Neuroscience, 17(5), 725–731. 10.1038/nn.3687

Moser, E., Moser, M. B., & Andersen, P. (1993). Spatial learning impairment parallels the magnitude of dorsal hippocampal lesions, but is hardly present following ventral lesions. Journal of Neuroscience, 13(9), 3916–3925. 10.1523/JNEUROSCI.13-09-03916.1993

Muller, R. U., & Kubie, J. L. (1987). The effects of changes in the environment on the spatial firing of hippocampal complex-spike cells. Journal of Neuroscience, 7(7), 1951–1968. 10.1523/JNEUROSCI.07-07-01951.1987

Nath, T., Mathis, A., Chen, A. C., Patel, A., Bethge, M., & Mathis, M. W. (2019). Using DeepLabCut for 3D markerless pose estimation across species and behaviors. Nature Protocols, 14(7), 2152–2176. 10.1038/s41596-019-0176-0

O’Keefe, J., & Conway, D. H. (1978). Hippocampal place units in the freely moving rat: Why they fire where they fire. Experimental Brain Research, 31(4), 573–590. 10.1007/BF00239813

O’Keefe, J., & Dostrovsky, J. (1971). The hippocampus as a spatial map: Preliminary evidence from unit activity in the freely-moving rat. Brain Research, 34, 171–175. 10.1016/0006-8993(71)90358-1

O’Keefe, J. & Nadel, L. (1978) *The Hippocampus as a Cognitive Map*. Oxford: Clarendon Press.

Packard, M. G., & McGaugh, J. L. (1996). Inactivation of Hippocampus or Caudate Nucleus with Lidocaine Differentially Affects Expression of Place and Response Learning. Neurobiology of Learning and Memory, 65(1), 65–72. 10.1006/nlme.1996.0007

Sargolini, F., Fyhn, M., Hafting, T., McNaughton, B. L., Witter, M. P., Moser, M.-B., & Moser, E. I. (2006). Conjunctive Representation of Position, Direction, and Velocity in Entorhinal Cortex. Science, 312(5774), 758–762. 10.1126/science.1125572

Scaplen, K. M., Gulati, A. A., Heimer-McGinn, V. L., & Burwell, R. D. (2014). Objects and landmarks: Hippocampal place cells respond differently to manipulations of visual cues depending on size, perspective, and experience. Hippocampus, 24(11), 1287–1299. 10.1002/hipo.22331

Sutherland, R. J., Whishaw, I. Q., & Kolb, B. (1983). A behavioural analysis of spatial localization following electrolytic, kainate- or colchicine-induced damage to the hippocampal formation in the rat. Behavioural Brain Research, 7(2), 133–153. 10.1016/0166-4328(83)90188-2

Sutherland, R. J., Whishaw, I. Q., & Regehr, J. C. (1982). Cholinergic receptor blockade impairs spatial localization by use of distal cues in the rat. 96(4), 563–573. 10.1037/h0077914

Taube, J. S. (2007). The Head Direction Signal: Origins and Sensory-Motor Integration. Annual Review of Neuroscience, 30(Volume 30, 2007), 181–207. 10.1146/annurev.neuro.29.051605.112854

Taube, J. S., Muller, R. U., & Ranck, J. B. (1990). Head-direction cells recorded from the postsubiculum in freely moving rats. I. Description and quantitative analysis. Journal of Neuroscience, 10(2), 420–435. 10.1523/JNEUROSCI.10-02-00420.1990

Tolman, E. C. (1948). Cognitive maps in rats and men. Psychological Review, 55, 189–208. 10.1037/h0061626

Vinogradova, O. S. (1995). Expression, control, and probable functional significance of the neuronal theta-rhythm. Progress in Neurobiology, 45(6), 523–583. 10.1016/0301-0082(94)00051-I

Wallace, D. G., Gorny, B., & Whishaw, I. Q. (2002). Rats can track odors, other rats, and themselves: Implications for the study of spatial behavior. Behavioural Brain Research, 131(1), 185–192. 10.1016/S0166-4328(01)00384-9

